# Selection and characterization of a reovirus mutant with improved thermostability

**DOI:** 10.1101/511352

**Authors:** Anthony J. Snyder, Pranav Danthi

## Abstract

The environment represents a significant barrier to infection. Physical stressors (heat) or chemical agents (ethanol and sodium dodecyl sulfate) can render virions noninfectious. As such, discrete proteins are necessary to stabilize the dual layered structure of mammalian orthoreovirus (reovirus). The outer capsid participates in cell entry: (i) σ3 is degraded to generate the infectious subviral particle and (ii) μ1 facilitates membrane penetration and subsequent core delivery. μ1-σ3 interactions also prevent inactivation; however, this activity is not fully characterized. Using forward and reverse genetic approaches, we identified two mutations (μ1 M258I and σ3 S344P) within heat resistant strains. σ3 S344P was sufficient to enhance capsid integrity and to reduce protease sensitivity. Moreover, these changes impaired replicative fitness in a reassortant background. This work reveals new details regarding the determinants of reovirus stability.

**SIGNIFICANCE:** Nonenveloped viruses rely on protein-protein interactions to shield their genomes from the environment. The capsid, or protective shell, must also disassemble during cell entry. In this work, we identified a determinant within mammalian orthoreovirus that regulates heat resistance, disassembly kinetics, and replicative fitness. Together, capsid function is balanced for optimal replication and for spread to a new host.

## INTRODUCTION

Mammalian orthoreovirus (reovirus) is a versatile system to explore the properties of dual layered structures (1). Reovirus is composed of two concentric, protein shells (2). Each component serves a structural and/or functional role in the replication cycle. The inner capsid (core) encapsidates 10 segments of genomic dsRNA (1) and supports polymerase activity during infection (3–5). The core is highly stable, presumably to shield the viral genome from host sensors (6). The outer capsid comprises 200 μ1-σ3 heterohexamers and a maximum of 12 σ1 trimers (2, 7) and is required for cell entry (1). Virions are differentially sensitive to inactivating agents; μ1 adopts an altered conformation at elevated temperatures (6).

Reovirus initiates infection by attaching to proteinaceous receptors (8–10) or serotype-specific glycans (11–14). Virions are internalized by receptor-mediated endocytosis (15–18) and traffic to late endosomes (19–23). Acid dependent cathepsin proteases degrade σ3 (24–28) and cleave μ1 into μ1δ and Φ (Fig. 2A) (29). The resulting intermediate is called the infectious subviral particle (30). Proteolytic disassembly (virion-to-ISVP conversion) is recapitulated *in vitro* by treating purified virions with exogenous protease (29–31). During subsequent rearrangements (ISVP-to-ISVP* conversion), neighboring μ1 trimers separate (7, 32) and μ1δ is cleaved into μ1N and δ (Fig. 2A) (33, 34). The δ fragment then adopts a hydrophobic and protease sensitive conformation (35). This step is accompanied by the release of μ1N and Φ pore forming peptides, which disrupt the endosomal membrane (33–40). ISVP-to-ISVP* conversion can be triggered *in vitro* using heat (35, 41).

Reovirus must remain environmentally stable prior to infection. Heat or chemical agents render particles noninfectious (42, 43). Nonetheless, rare subpopulations harbor resistance-granting mutations, likely due to error prone replication. Studies of such variants provide clues regarding capsid stability and/or structure. When comparing prototype strains, differences in the efficiency of inactivation map to either the M2 gene segment (encodes for μ1) or the S4 gene segment (encodes for σ3) (35, 41, 43). μ1 mutations were selected by exposing virions to ethanol or by exposing ISVPs to heat. These changes reveal important μ1 mediated intratrimer, intertrimer, and trimer-core contacts (44–47). Similarly, σ3 Y354H was selected from persistently infected cells. This change alters capsid properties and allows for disassembly under limiting cathepsin activity (48, 49). σ3 also represents the primary determinant for virion thermostability (43). μ1-σ3 interactions are thought to prevent irreversible and premature, entry related conformational changes (6); however, this idea has not been fully investigated. In this work, we selected for and characterized heat resistant (HR) strains: (i) μ1 M258I and σ3 S344P were identified within HR strains, (ii) σ3 S344P was sufficient to enhance capsid integrity and to reduce protease sensitivity, and (iii) HR mutations impaired replicative fitness in a reassortant background.

## MATERIALS AND METHODS

### Cells and viruses

Murine L929 (L) cells were grown at 37°C in Joklik’s minimal essential medium (Lonza) supplemented with 5% fetal bovine serum (Life Technologies), 2 mM L-glutamine (Invitrogen), 100 U/ml penicillin (Invitrogen), 100 μg/ml streptomycin (Invitrogen), and 25 ng/ml amphotericin B (Sigma-Aldrich). All viruses used in this study were derived from reovirus type 1 Lang (T1L) and reovirus type 3 Dearing (T3D) and were generated by plasmid-based reverse genetics (50, 51). Mutations within the T1L S4, T1L M2, and T3D M2 genes were generated by QuikChange site-directed mutagenesis (Agilent Technologies). S344P in T1L σ3 was made using the following primer pair: forward 5’-CTGCTCTCACAATGTTCCCGGACACCACCAAGTTCGG-3’ and reverse 5’-CCGAACTTGGTGGTGTCCGGGAACATTGTGAGAGCAG-3’. M258I in T1L μ1 was made using the following primer pair: forward 5’-TCAGAAGGAACTGTGATTAATGAGGCCGTGAATGC-3’ and reverse 5’-GCATTCACGGCCTCATTAATCACAGTTCCTTCTGA-3’. M258I in T3D μ1 was made using the following primer pair: forward 5’-ACGTATCAGAAGGCACCGTGATTAACGAGGCTGTC-3’ and reverse 5’-GACAGCCTCGTTAATCACGGTGCCTTCTGATACGT-3’.

### Virion purification

Recombinant reoviruses T1L, T1L/T3D M2, T1L μ1 M258I, T1L σ3 S344P, T1L μ1 M258I σ3 S344P, and T1L/T3D M2 μ1 M258I σ3 S344P were propagated and purified as previously described (50, 51). All variants in the T1L/T3D M2 background contained a T3D M2 gene in an otherwise T1L background. L cells infected with second passage reovirus stocks were lysed by sonication. Virions were extracted from lysates using Vertrel-XF specialty fluid (Dupont) (52). The extracted particles were layered onto 1.2-to 1.4-g/cm^3^ CsCl step gradients. The gradients were then centrifuged at 187,000×*g* for 4 h at 4°C. Bands corresponding to purified virions (~1.36 g/cm^3^) (53) were isolated and dialyzed into virus storage buffer (10 mM Tris, pH 7.4, 15 mM MgCl_2_, and 150 mM NaCl). Following dialysis, the particle concentration was determined by measuring the optical density of the purified virion stocks at 260 nm (OD_260_; 1 unit at OD_260_ = 2.1 ×10^12^ particles/ml) (54). The purification of virions was confirmed by SDS-PAGE and Coomassie brilliant blue (Sigma-Aldrich) staining.

### Selection and isolation of heat resistant (HR) strains

T1L virions (2×10^12^ particles/ml) were incubated for 5 min at 55°C in a S1000 thermal cycler (Bio-Rad). The total volume of the reaction was 30 μl in virus storage buffer (10 mM Tris, pH 7.4, 15 mM MgCl_2_, and 150 mM NaCl). Following incubation, 10 μl were analyzed by plaque assay. Putative HR strains were plaque purified at 5 days post infection. Next, L cell monolayers in 60 mm dishes (Greiner Bio-One) were adsorbed with the isolated plaques for 1 h at 4°C. Following the viral attachment incubation, the monolayers were washed three times with ice-cold PBS and overlaid with 2 ml of Joklik’s minimal essential medium (Lonza) supplemented with 5% fetal bovine serum (Life Technologies), 2 mM L-glutamine (Invitrogen), 100 U/ml penicillin (Invitrogen), 100 μg/ml streptomycin (Invitrogen), and 25 ng/ml amphotericin B (Sigma-Aldrich). The cells were incubated at 37°C until cytopathic effect was observed and then lysed by two freeze-thaw cycles (first passage). To verify heat resistance, the infected cell lysates were incubated for 5 min at 55°C in a S1000 thermal cycler (Bio-Rad). The total volume of each reaction was 30 μl. For each isolate, an aliquot was also incubated for 5 min at 4°C. Following incubation, 10 μl of each reaction were diluted into 40 μl of ice-cold virus storage buffer (10 mM Tris, pH 7.4, 15 mM MgCl_2_, and 150 mM NaCl) and infectivity was determined by plaque assay. The change in infectivity was calculated using the following formula: log_10_(PFU/ml)_55°c_ -log_10_(PFU/ml)_4°c_.

### Sequencing of HR strains

L cell monolayers in 60 mm dishes (Greiner Bio-One) were adsorbed with first passage, verified HR strains for 1 h at 4°C. Following the viral attachment incubation, the monolayers were washed three times with ice-cold PBS and overlaid with 2 ml of Joklik’s minimal essential medium (Lonza) supplemented with 5% fetal bovine serum (Life Technologies), 2 mM L-glutamine (Invitrogen), 100 U/ml penicillin (Invitrogen), 100 μg/ml streptomycin (Invitrogen), and 25 ng/ml amphotericin B (Sigma-Aldrich). At 24 h post infection, the cells were lysed with TRI Reagent (Molecular Research Center). Viral RNA was isolated by phenol-chloroform extraction and then subjected to reverse transcription (RT)-PCR using T1L S4 or T1L M2 gene segment-specific primers. PCR products were resolved on Tris-acetate-EDTA agarose gels, purified using a QIAquick gel extraction kit (Qiagen), and sequenced. The identified mutations were reintroduced into clean, T1L and T1L/T3D M2 backgrounds.

### Sequence analysis

Multiple sequence alignments were created using the Clustal Omega program (55).

### Structure analysis

Molecular graphics were created using the UCSF Chimera program (56).

### Generation of infectious subviral particles (ISVPs)

T1L, T1L/T3D M2, T1L μ1 M258I, T1L σ3 S344P, T1L μ1 M258I σ3 S344P, or T1L/T3D M2 μ1 M258I σ3 S344P virions (2×10^12^ particles/ml) were digested with 200 μg/ml TLCK (*N*α-*p*-tosyl-L-lysine chloromethyl ketone)-treated chymotrypsin (Worthington Biochemical) in a total volume of 100 μl for 20 min at 32°C (30, 31). The reactions were then incubated on ice for 20 min and quenched by the addition of 1 mM phenylmethylsulfonyl fluoride (PMSF) (Sigma-Aldrich). The generation of ISVPs was confirmed by SDS-PAGE and Coomassie brilliant blue (Sigma-Aldrich) staining.

### Dynamic light scattering

T1L, T1L/T3D M2, T1L μ1 M258I, T1L σ3 S344P, T1L μ1 M258I σ3 S344P, or T1L/T3D M2 μ1 M258I σ3 S344P virions or ISVPs (2×10^12^ particles/ml) were analyzed using a Zetasizer Nano S dynamic light scattering system (Malvern Instruments). All measurements were made at room temperature in a quartz Suprasil cuvette with a 3.00-mm-path length (Hellma Analytics). For each sample, the size distribution profile was determined by averaging readings across 15 iterations.

### Thermal inactivation and trypsin sensitivity assays

T1L, T1L/T3D M2, T1L μ1 M258I, T1L σ3 S344P, T1L μ1 M258I σ3 S344P, or T1L/T3D M2 μ1 M258I σ3 S344P virions or ISVPs (2×10^12^ particles/ml) were incubated for 5 min at the indicated temperatures in a S1000 thermal cycler (Bio-Rad). The total volume of each reaction was 30 μl in virus storage buffer (10 mM Tris, pH 7.4, 15 mM MgCl_2_, and 150 mM NaCl). For each reaction condition, an aliquot was also incubated for 5 min at 4°C. Following incubation, 10 μl of each reaction were diluted into 40 μl of ice-cold virus storage buffer (10 mM Tris, pH 7.4, 15 mM MgCl_2_, and 150 mM NaCl) and infectivity was determined by plaque assay. The change in infectivity at a given temperature (T) was calculated using the following formula: log_10_(PFU/ml)_*T*_ -log_10_(PFU/ml)_4°C_. The titers of the 4°C control samples were between 5×10^9^ and 5×10^10^ PFU/ml. The remaining 20 μl of each reaction were either mock treated or treated with 0.08 mg/ml trypsin (Sigma-Aldrich) for 30 min on ice. Following digestion, equal particle numbers from each reaction were solubilized in reducing SDS sample buffer and analyzed by SDS-PAGE. The gels were Coomassie Brilliant Blue (Sigma-Aldrich) stained and imaged on an Odyssey imaging system (LI-COR).

### Degradation of σ3 by exogenous protease

T1L, T1L/T3D M2, T1L μ1 M258I, T1L σ3 S344P, T1L μ1 M258I σ3 S344P, or T1L/T3D M2 μ1 M258I σ3 S344P virions (2×10^12^ particles/ml) were incubated in the presence of 10 μg/ml endoproteinase LysC (New England Biolabs) at 37°C in a S1000 thermal cycler (Bio-Rad). The starting volume of each reaction was 100 μl in virus storage buffer (10 mM Tris, pH 7.4, 15 mM MgCl_2_, and 150 mM NaCl). At the indicated time points, 10 μl of each reaction was solubilized in reducing SDS sample buffer and boiled for 10 min at 95°C. Equal particle numbers from each time point were analyzed by SDS-PAGE. The gels were Coomassie Brilliant Blue (Sigma-Aldrich) stained and imaged on an Odyssey imaging system (LI-COR).

### Ammonium chloride (AC) escape assay

L cell monolayers in 6-well plates (Greiner Bio-One) were adsorbed with T1L, T1L μ1 M258I, T1L σ3 S344P, or T1L μ1 M258I σ3 S344P virions (10 PFU/cell) for 1 h at 4°C. Following the viral attachment incubation, the monolayers were washed three times with ice-cold PBS and overlaid with 2 ml of Joklik’s minimal essential medium (Lonza) supplemented with 5% fetal bovine serum (Life Technologies), 2 mM L-glutamine (Invitrogen), 100 U/ml penicillin (Invitrogen), 100 μg/ml streptomycin (Invitrogen), and 25 ng/ml amphotericin B (Sigma-Aldrich). The cells were either lysed immediately by two freeze-thaw cycles (input) or incubated at 37°C (start of infection). At the indicated times post infection, the growth medium was supplemented with 20 mM AC (Mallinckrodt Pharmaceuticals). At 24 h post infection, the cells were lysed by two freeze-thaw cycles and the virus titer was determined by plaque assay. The viral yield for each infection condition (timing of AC addition) (t) was calculated using the following formula: log_10_(PFU/ml)_*t*_ -log_10_(PFU/ml)_input_. The titers of the input samples were between 1 ×10^5^ and 4×10^5^ PFU/ml.

### Single- and multistep growth assays

L cell monolayers in 6-well plates (Greiner Bio-One) were adsorbed with T1L, T1L/T3D M2, T1L μ1 M258I, T1L σ3 S344P, T1L μ1 M258I σ3 S344P, or T1L/T3D M2 μ1 M258I σ3 S344P virions (10 PFU/cell or 0.01 PFU/cell) for 1 h at 4°C. Following the viral attachment incubation, the monolayers were washed three times with ice-cold PBS and overlaid with 2 ml of Joklik’s minimal essential medium (Lonza) supplemented with 5% fetal bovine serum (Life Technologies), 2 mM L-glutamine (Invitrogen), 100 U/ml penicillin (Invitrogen), 100 μg/ml streptomycin (Invitrogen), and 25 ng/ml amphotericin B (Sigma-Aldrich). The cells were either lysed immediately by two freeze-thaw cycles (input) or incubated at 37°C (start of infection). At the indicated times post infection, the cells were lysed by two freeze-thaw cycles and the virus titer was determined by plaque assay. The viral yield at a given time post infection (*t*) was calculated using the following formula: log_10_(PFU/ml)_*t*_ -log_10_(PFU/ml)_input_. Following infection with 10 PFU/cell, the titers of the input samples were between 1×10^5^ and 4×10^5^ PFU/ml. Following infection with 0.01 PFU/cell, the titers of the input samples were between 1×10^2^ and 3×10^2^ PFU/ml.

### Plaque assay

Control or heat-treated virus samples or infected cell lysates were diluted in PBS supplemented with 2 mM MgCl_2_ (PBS^Mg^). L cell monolayers in 6-well plates (Greiner Bio-One) were infected with 250 μl of diluted virus for 1 h at room temperature. Following the viral attachment incubation, the monolayers were overlaid with 4 ml of serum-free medium 199 (Sigma-Aldrich) supplemented with 1% Bacto Agar (BD Biosciences), 10 μg/ml TLCK-treated chymotrypsin (Worthington Biochemical), 2 mM L-glutamine (Invitrogen), 100 U/ml penicillin (Invitrogen), 100 μg/ml streptomycin (Invitrogen), and 25 ng/ml amphotericin B (Sigma-Aldrich). The infected cells were incubated at 37°C, and plaques were counted at 5 days post infection.

### Statistical analysis

Unless noted otherwise, the reported values represent the mean of three independent, biological replicates. Error bars indicate standard deviation. *P* values were calculated using Student’s *t* test (two-tailed, unequal variance assumed). For thermal inactivation experiments (Figs. 1A, 3A, and 5C), two criteria were used to assign significance: *P* ≤ 0.05 and difference in change in infectivity ≥ 2 log_10_ units. For single- and multistep growth experiments (Fig. 6), two criteria were used to assign significance: *P* ≤ 0.05 and difference in viral yield ≥ 1 log_10_ unit.

## RESULTS AND DISCUSSION

### Characterization of putative heat resistant (HR) strains

Reovirus is susceptible to environmental factors (42, 43). The loss of infectivity is correlated with outer capsid rearrangements (6, 35, 41). To isolate HR strains, type 1 Lang (T1L) virions were incubated at 55°C. The virus titer decreased by ~5.5 log10 units compared to the input (data not shown). Survivors were plaque purified and used to generate infected cell lysates. Three plaque isolates (out of 5) exhibited a bona fide HR phenotype (Fig. 1A). Sequencing of the μ1- and σ3-encoding gene segments (M2 and S4, respectively) revealed two mutations within each HR strain: μ1 M258I (conserved position) and σ3 S344P (nonconserved position) (Fig. 1B). These residues, which are not expected to interact, occupy distinct positions within the T1L μ1-σ3 heterohexamer (Figs. 1C-D) (7).

**Fig 1.**
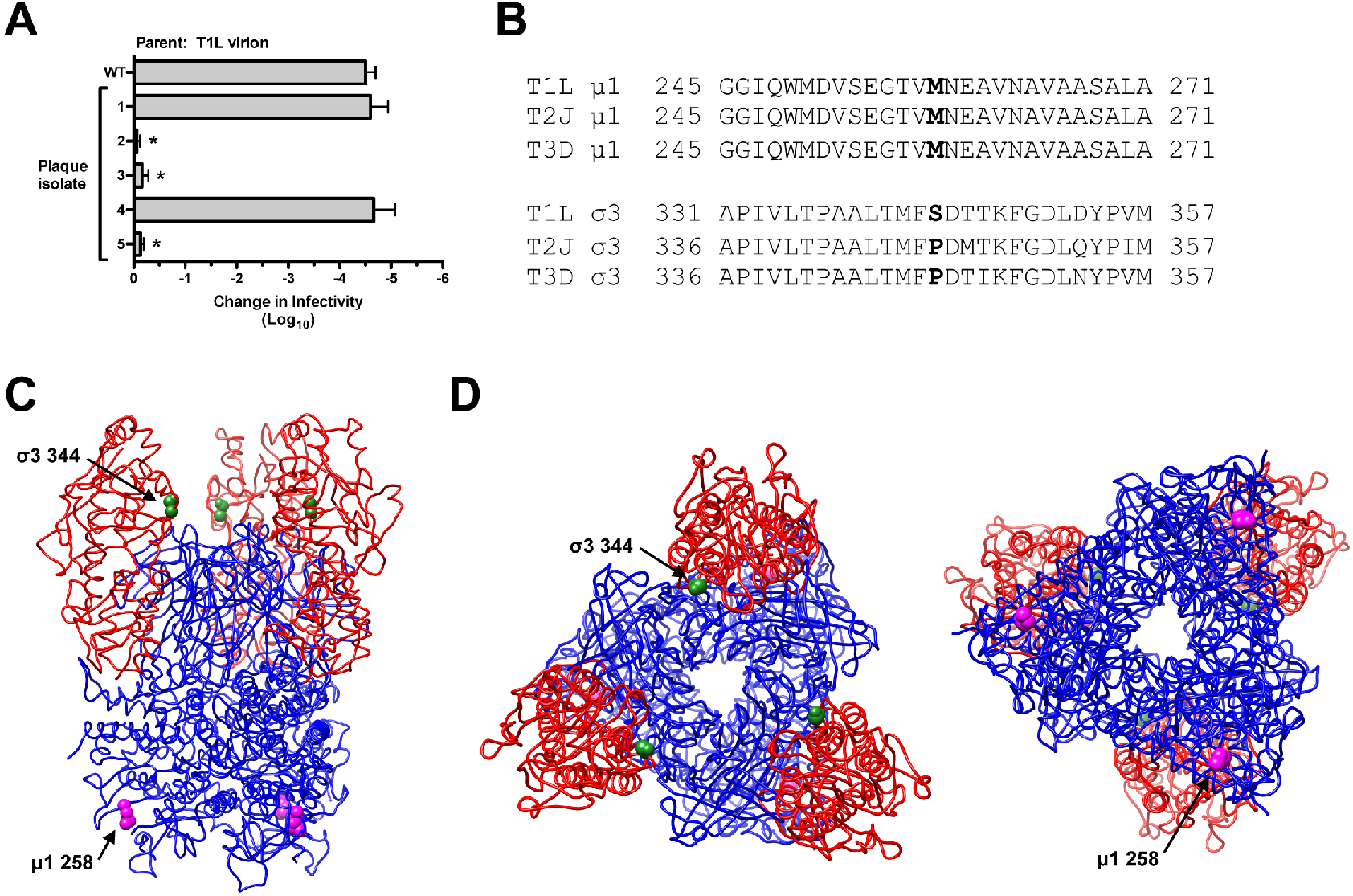
Selection, isolation, and sequencing of heat resistant strains. (A) Thermal inactivation. T1L virions were incubated for 5 min at 55^8^C. Surviving strains were plaque purified and amplified on L cells. To verify heat resistance, the infected cell lysates were incubated for 5 min at 55°C. The change in infectivity relative to samples incubated at 4°C was determined by plaque assay. The data are presented as means ± SDs. *, *P*≤ 0.05 and difference in change in infectivity ≥ 2 log_10_ units (n = 3 independent replicates). (B) Multiple sequence alignments. Residues corresponding to μ1 258 and σ3 344 are in boldface. (C) Side view of the T1L μ1-σ3 heterohexamer (7) (Protein Data Bank [PDB] accession number 1JMU). (D) Top and bottom views (left and right, respectively) of the T1L μ1-σ3 heterohexamer (7) (Protein Data Bank [PDB] accession number 1JMU). In panels C and D, μ1 monomers are colored blue, σ3 monomers are colored red, μ1 258 is represented by magenta spheres, and σ3 344 is represented by green spheres.

### σ3 S344P was sufficient to enhance capsid integrity

Changes within other capsid components (σ1 and core proteins) could influence the HR phenotype. To address this concern, we reintroduced μ1 M258I and σ3 S344P into clean T1L backgrounds. Variants with one or both mutations displayed no observable defects in protein composition or protein stoichiometry. Due to autocatalytic activity, μ1 resolves as μ1C, and μ1δ resolves as δ (Figs. 2A-B) (33). We also analyzed virions and ISVPs by dynamic light scattering (DLS). In each case, we detected a single peak with no evidence of aggregation (Fig. 2C).

**Fig 2.**
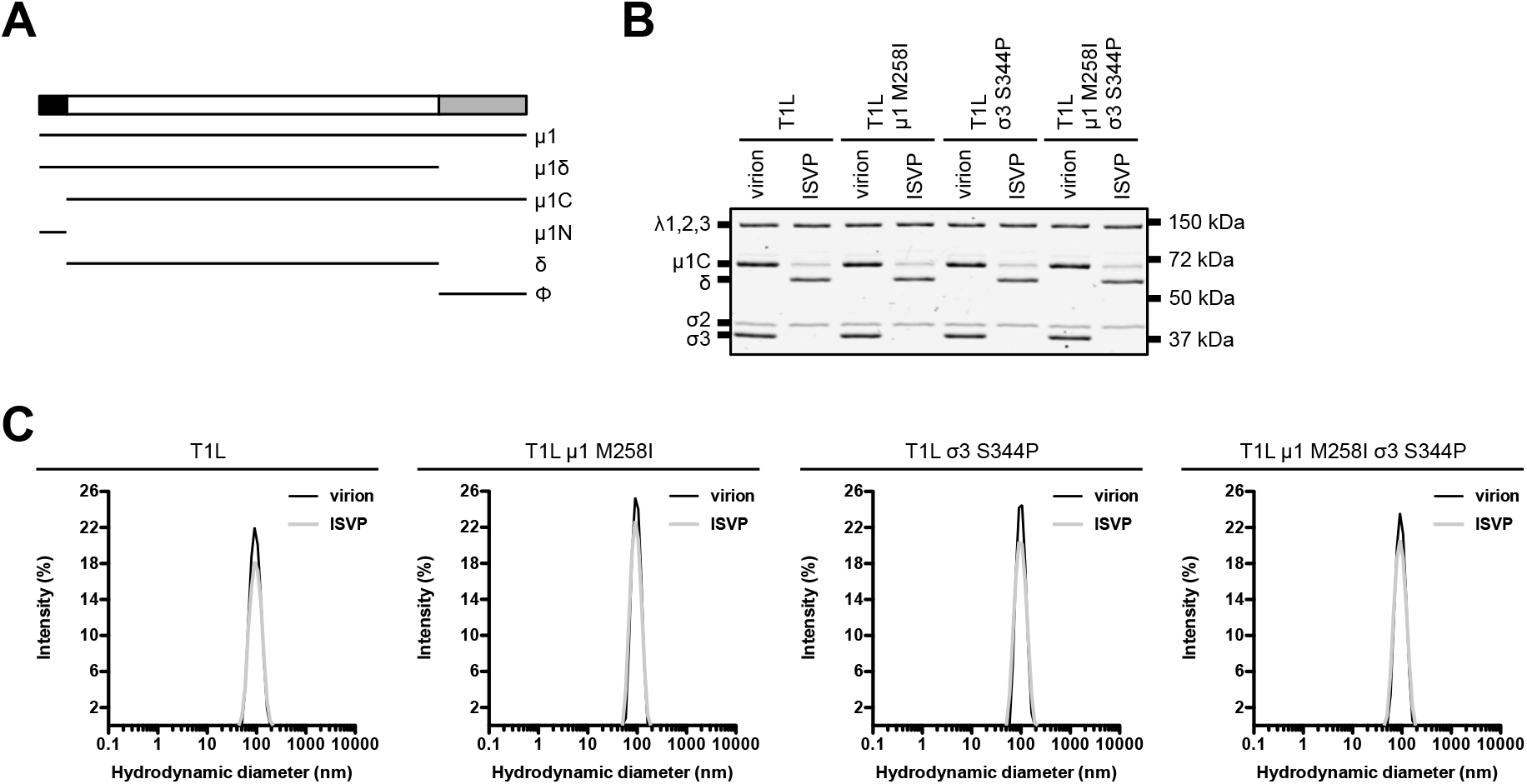
Protein compositions and size distribution profiles of T1L variants. (A) Schematic of μ1 cleavage fragments. (B) Protein compositions. T1L, T1L μ1 M258I, T1L σ3 S344P, and T1L μ1 M258I σ3 S344P virions and ISVPs were analyzed by SDS-PAGE. The gel was Coomassie brilliant blue stained. The migration of capsid proteins is indicated on the left. μ1 resolves as μ1C, and μ1δ resolves as δ (33). μ1N and Φ are too small to resolve on the gel (n = 3 independent replicates; results from 1 representative experiment are shown). (C) Size distribution profiles. T1L, T1L μ1 M258I, T1L σ3 S344P, or T1L μ1 M258I σ3 S344P virions or ISVPs were analyzed by dynamic light scattering. For each variant, the virion (black) and ISVP (gray) size distribution profiles are overlaid (n = 3 independent replicates; results from 1 representative experiment are shown).

The S4 gene segment (encodes for σ3) contains the genetic determinants for virion thermostability (43). σ3 preserves infectivity by stabilizing μ1 (6). Any change that affects μ1-σ3 structure could modulate this activity. To test this idea, we performed thermal inactivation experiments. Following incubation at 55°C, T1L and T1L μ1 M258I virions were reduced in titer by ~5.5 log_10_ units relative to control that was incubated at 4°C. In contrast, T1L σ3 S344P and T1L μ1 M258I σ3 S344P virions were reduced in titer by ~0.5 log_10_ units after incubation at 55°C and by ~4.0 log_10_ units after incubation at 58°C (Fig. 3A). Virion associated μ1 adopts an ISVP*-like (protease sensitive) conformation concurrent with inactivation. This transition is assayed *in vitro* by determining the susceptibility of μ1 to trypsin digestion (6, 35, 41). Consistent with the above results, 55°C was the minimal temperature at which μ1 in T1L and T1L μ1 M258I virions became trypsin sensitive, whereas 58°C was the minimal temperature at which μ1 in T1L σ3 S344P and T1L μ1 M258I σ3 S344P virions became trypsin sensitive (Fig. 3B). Of note, σ3 was absent from gels and μ1 migrated as uncleaved μ1C and cleaved δ. Trypsin, which was used to probe for protease sensitivity, degrades σ3 and cleaves at the μ1 δ-Φ junction (29). Heating alone was not sufficient to alter μ1 or σ3 levels (Fig. 3C).

**Fig 3.**
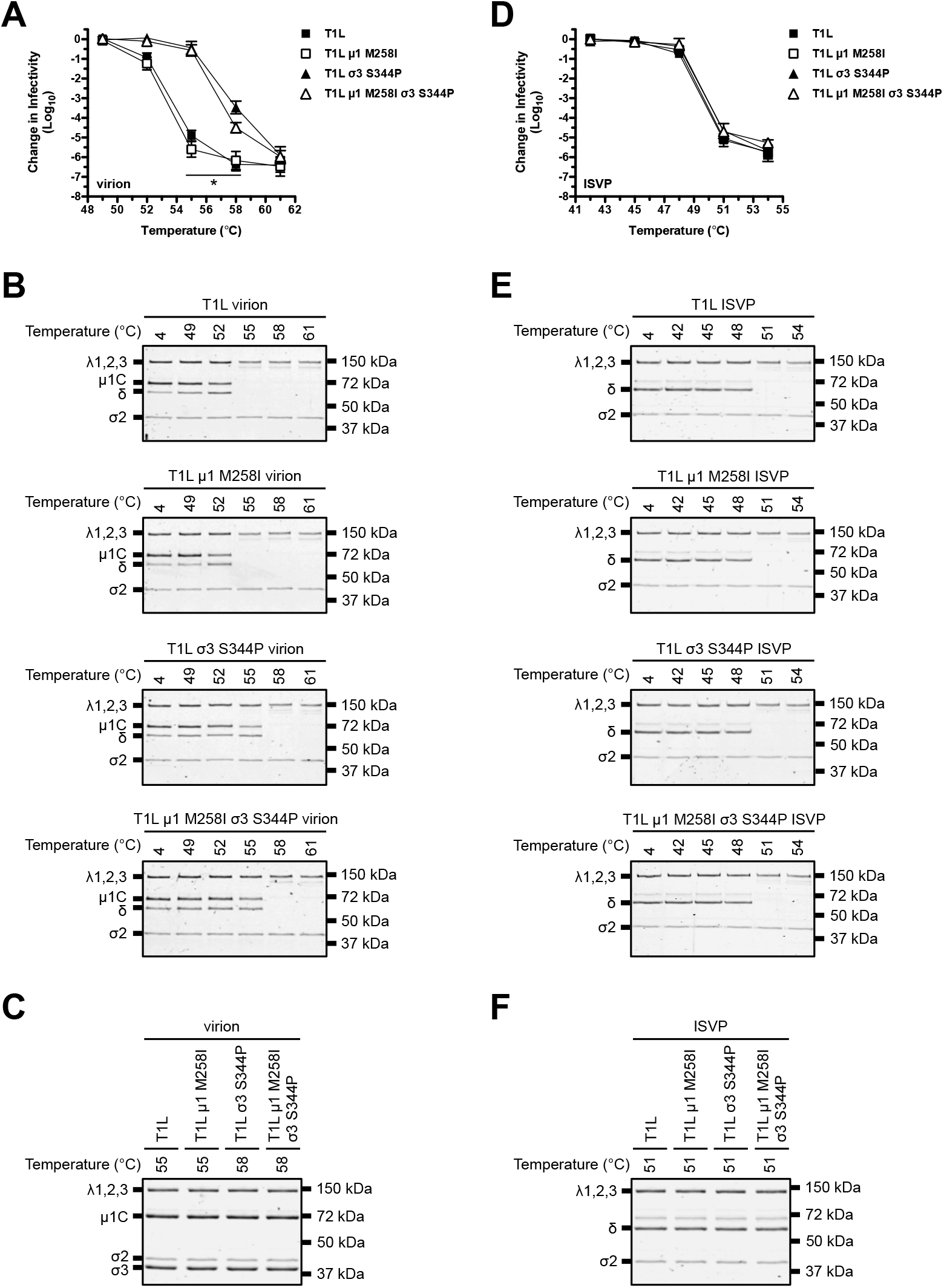
Thermostability of T1L variants. (A and D) Thermal inactivation. T1L, T1L μ1 M258I, T1L σ3 S344P, or T1L μ1 M258I σ3 S344P virions (A) or ISVPs (D) were incubated in virus storage for 5 min at the indicated temperatures. The change in infectivity relative to samples incubated at 4°C was determined by plaque assay. The data are presented as means ± SDs. *, *P* ≤ 0.05 and difference in change in infectivity ≥ 2 log_10_ units (n = 3 independent replicates). (B and E) Heat induced conformational changes. T1L, T1L μ1 M258I, T1L σ3 S344P, or T1L μ1 M258I σ3 S344P virions (B) or ISVPs (E) were incubated in virus storage buffer for 5 min at the indicated temperatures. Each reaction was then treated with trypsin for 30 min on ice. Following digestion, equal particle numbers from each reaction were analyzed by SDS-PAGE. The gels were Coomassie brilliant blue stained (n = 3 independent replicates; results from 1 representative experiment are shown). (C and F). Composition of heated virus. T1L, T1L μ1 M258I, T1L σ3 S344P, or T1L μ1 M258I σ3 S344P virions (C) or ISVPs (F) were incubated in virus storage buffer for 5 min at the indicated temperatures. Equal particle numbers from each reaction were analyzed by SDS-PAGE. The gels were Coomassie brilliant blue stained (n = 3 independent replicates; results from 1 representative experiment are shown).

The M2 gene segment (encodes for μ1) contains the genetic determinants for ISVP thermostability. The transition to ISVP* induces the loss of infectivity (35, 41). Following incubation at 51°C, T1L, T1L μ1 M258I, T1L σ3 S344P, and T1L μ1 M258I σ3 S344P ISVPs were reduced in titer by ~5.0 log_10_ units (Fig. 3D). Thus, HR mutations conferred stability only within the context of a virion (σ3 degradation restored wild-type-like heat sensitivity). Protease treatment also serves as a biochemical probe for ISVP* formation (35, 41). For each variant, δ (a product of μ1 cleavage) (Fig. 2A) became trypsin sensitive at 51°C (Figs 3E). Heating alone was not sufficient to induce the loss of δ (Fig. 3F).

### σ3 S344P was sufficient to reduce protease sensitivity

Proteolytic disassembly (virion-to-ISVP conversion) is required for reovirus to establish an infection (27). T1L is more protease sensitive than T3D. This difference was mapped to polymorphisms at 344, 347, and 353 using recombinant protein (57). σ3 354 also regulates capsid properties (48, 49). These residues are thought to influence conformational flexibility through specific, intramonomer contacts (7, 57, 58). To determine if HR mutations alter disassembly kinetics, virions were digested *in vitro* with endoproteinase LysC (EKC). EKC probes for subtle differences in structure (57, 59). T1L and T1L μ1 M258I σ3 were degraded within 40 min. In contrast, T1L σ3 S344P and T1L μ1 M258I σ3 S344P σ3 persisted (in part) for 100 min (Fig. 4A). We next tested the sensitivity to intracellular proteases. Cathepsin B-L require endosomal acidification for activity. As such, lysosomotropic weak bases (ammonium chloride [AC]) block infection (27, 60). Murine L929 (L) cells were adsorbed with virions, and viral yield was quantified at 24 h post infection. When indicated, the growth medium was supplemented with AC. The timing of AC escape is related to the rate of disassembly (27). Consistent with the above results, T1L and T1L μ1 M258I bypassed the block to infection with faster kinetics (viral yield of ~1.5 log_10_ units when AC was added at 60 min) than T1L σ3 S344P and T1L μ1 M258I σ3 S344P (viral yield of ~1.5 log_10_ units when AC was added at 90 min) (Fig. 4B). These results provide evidence that HR mutations diminish conformational flexibility; enhanced capsid integrity (Fig. 3) was correlated with reduced protease sensitivity (Fig. 4). σ3 is initially cleaved in a hypersensitive region between 208-214 or 238-250 (57). Presumably, σ3 S344P influenced protease accessibility. This idea was previously suggested for σ3 Y354H (49). Of note, we did not observe a phenotype for μ1 M258I. This change was identified within each HR strain; however, its function was not determined.

**Fig 4.**
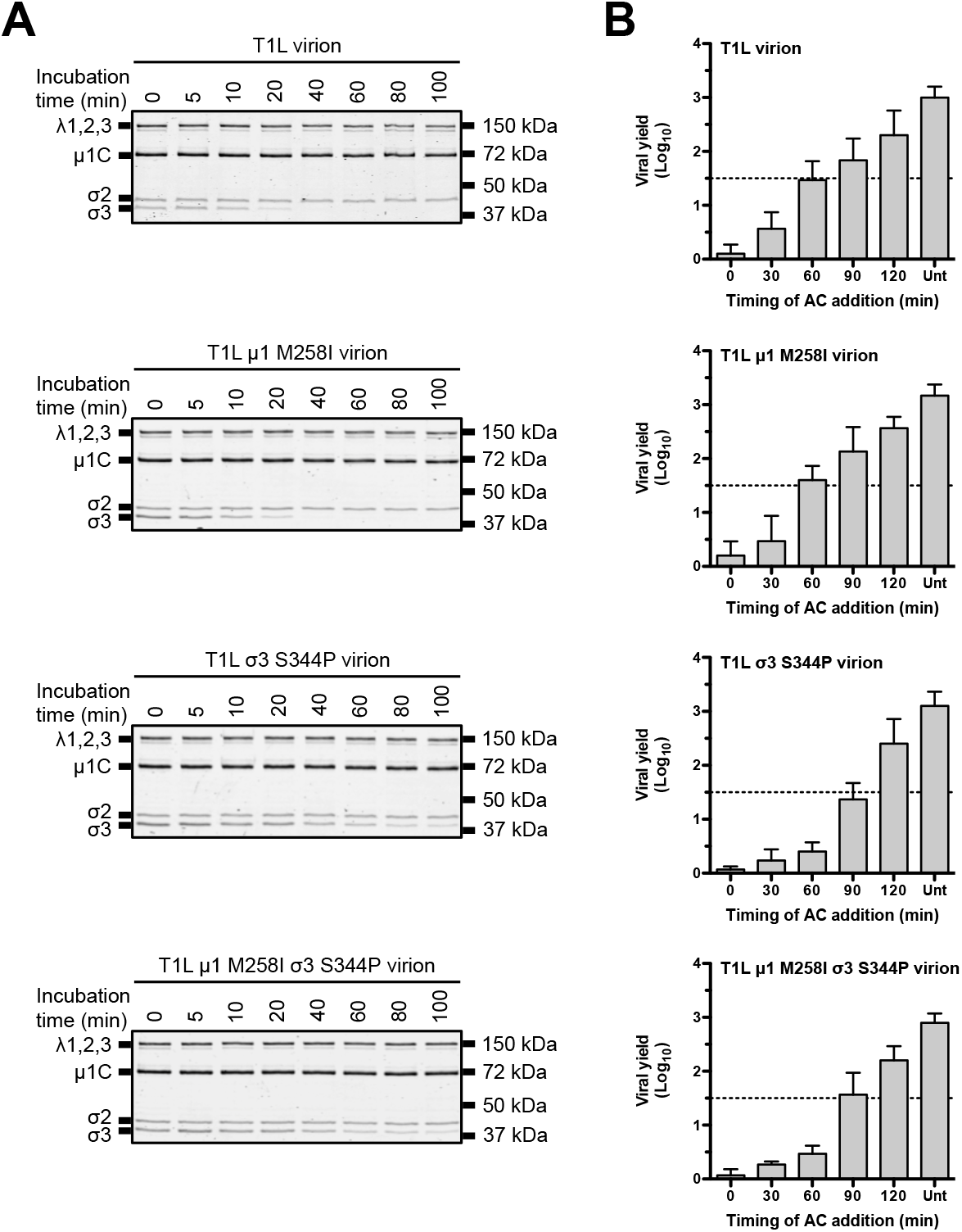
Degradation of σ3 by exogenous and intracellular proteases. (A) Exogenous protease. T1L, T1L μ1 M258I, T1L σ3 S344P, or T1L μ1 M258I σ3 S344P virions were incubated in virus storage buffer supplemented with endoproteinase LysC for the indicated amounts of time at 37°C. Following digestion, equal particle numbers from each time point were analyzed by SDS-PAGE. The gels were Coomassie brilliant blue stained (n = 3 independent replicates; results from 1 representative experiment are shown). (B) Intracellular proteases. L cell monolayers were infected with T1L, T1L μ1 M258I, T1L σ3 S344P, or T1L μ1 M258I σ3 S344P virions. At the indicated times post infection, the growth medium was supplemented with ammonium chloride. At 24 h post infection, the cells were lysed and viral yield was quantified by plaque assay. The data are presented as means ± SDs (n = 3 independent replicates). AC, ammonium chloride; Unt, untreated.

### HR mutations altered capsid properties in a reassortant background

T1L×T3D reassortants are used extensively to study many aspects of the viral replication cycle (1). For example, T1L/T3D M2 contains mismatched subunits (μ1-σ3 and μ1-core are derived from different strains). Virions with imperfect interactions retain wild-type-like stability and structure (6); however, HR mutations could function in a background dependent manner. To address this question, we generated T1L/T3D M2 μ1 M258I σ3 S344P. This virus displayed no observable defects in protein composition, protein stoichiometry, or particle size distribution (Figs. 5A-B). We next tested the impact on capsid properties. Following incubation at 55°C, T1L/T3D M2 virions were reduced in titer by ~5.5 log_10_ units, whereas T1L/T3D M2 μ1 M258I σ3 S344P virions were reduced in titer by ~1.0 log_10_ unit. In contrast, ISVPs were equally thermostable (Fig. 5C). HR mutations also conferred differential sensitivity to EKC. T1L/T3D M2 σ3 was degraded within 40 min, whereas T1L/T3D M2 μ1 M258I σ3 S344P σ3 persisted (in part) for 100 min (Fig. 5D).

**Fig 5.**
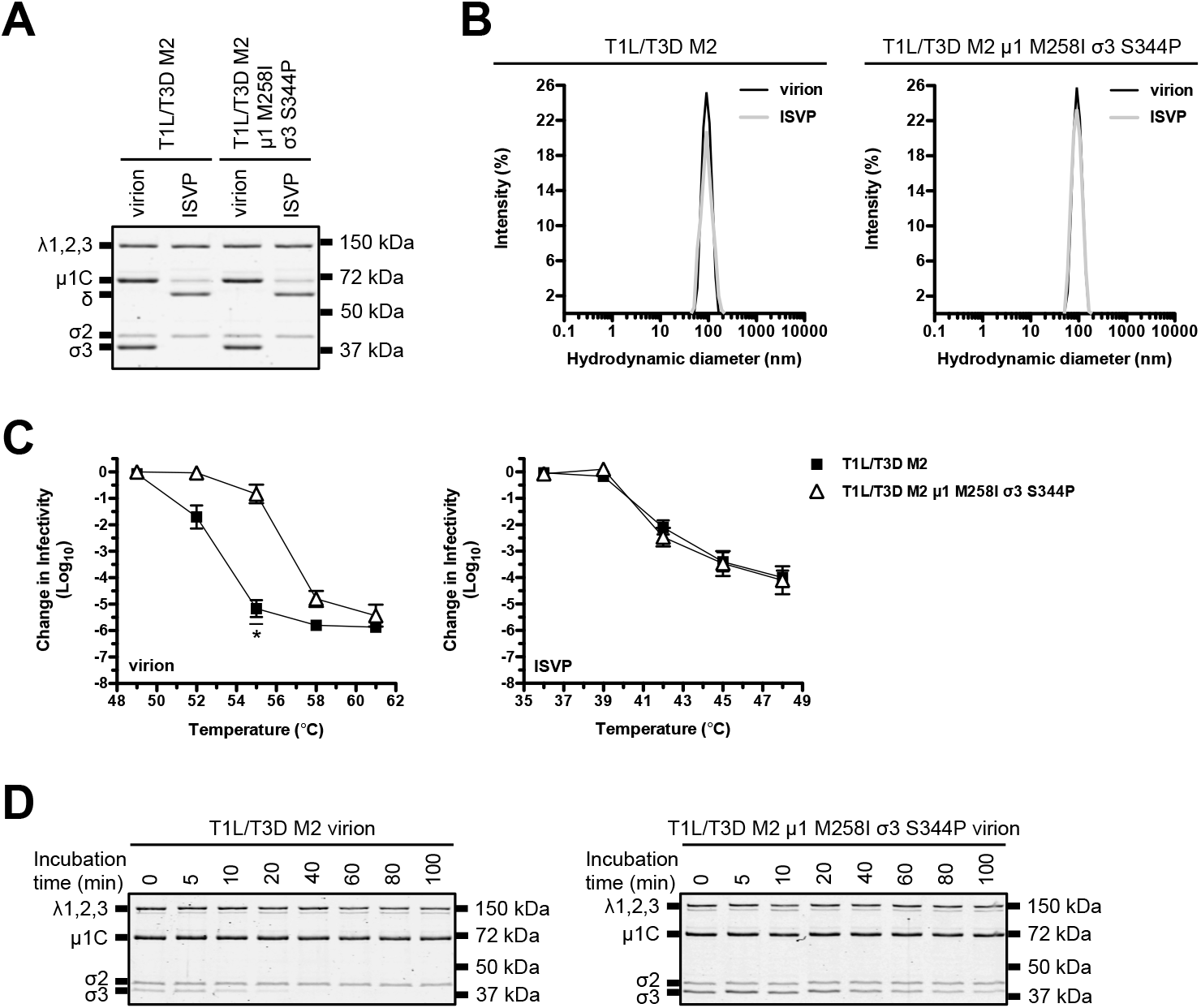
Thermostability of a T1L/T3D M2 variant. (A) Protein compositions. T1L/T3D M2 and T1L/T3D M2 μ1 M258I σ3 S344P virions and ISVPs were analyzed by SDS-PAGE. The gel was Coomassie brilliant blue stained. The migration of capsid proteins is indicated on the left. μ1 resolves as μ1C, and μ1δ resolves as δ (33). μ1N and Φ are too small to resolve on the gel (n = 3 independent replicates; results from 1 representative experiment are shown). (B) Size distribution profiles. T1L/T3D M2 or T1L/T3D M2 μ1 M258I σ3 S344P virions or ISVPs were analyzed by dynamic light scattering. For each variant, the virion (black) and ISVP (gray) size distribution profiles are overlaid (n = 3 independent replicates; results from 1 representative experiment are shown). (C) Thermal inactivation. T1L/T3D M2 or T1L/T3D M2 μ1 M258I σ3 S344P virions or ISVPs were incubated in virus storage for 5 min at the indicated temperatures. The change in infectivity relative to samples incubated at 4°C was determined by plaque assay. The data are presented as means ± SDs. *, *P*≤ 0.05 and difference in change in infectivity ≥ 2 log_10_ units (n = 3 independent replicates). (D) Degradation of σ3 by exogenous protease. T1L/T3D M2 or T1L/T3D M2 μ1 M258I σ3 S344P virions were incubated in virus storage buffer supplemented with endoproteinase LysC for the indicated amounts of time at 37°C. Following digestion, equal particle numbers from each time point were analyzed by SDS-PAGE. The gels were Coomassie brilliant blue stained (n = 3 independent replicates; results from 1 representative experiment are shown).

### HR mutations impaired replicative fitness in a reassortant background

Reovirus disassembles efficiently during cell entry, yet remains stable in the environment. This balance is necessary for a productive infection and for spread to a new host (1, 61). HR mutations are favored at high temperatures (Figs. 3 and 5). We next examined their impact under physiological conditions. L cells were infected at high (10 PFU/cell) or low (0.01 PFU/cell) MOI, and viral yield was quantified at the indicated times post infection. Each T1L variant grew to similar levels (Fig. 6A). Thus, biochemical differences (described above) do not confer a selective advantage (or impediment) during a bona fide infection. In contrast, T1L/T3D M2 μ1 M258I σ3 S344P produced fewer infectious units than T1L/T3D M2 by 24 h post infection (high MOI) and by 48 and 72 h post infection (low MOI) (Fig. 6B). Interestingly, we attempted to isolate unique, HR strains in the reassortant background; however, each plaque isolate failed secondary screening (data not shown). The reduced viral yield and the absence (or low abundance) of resistant strains imply a replication defect.

**Fig 6.**
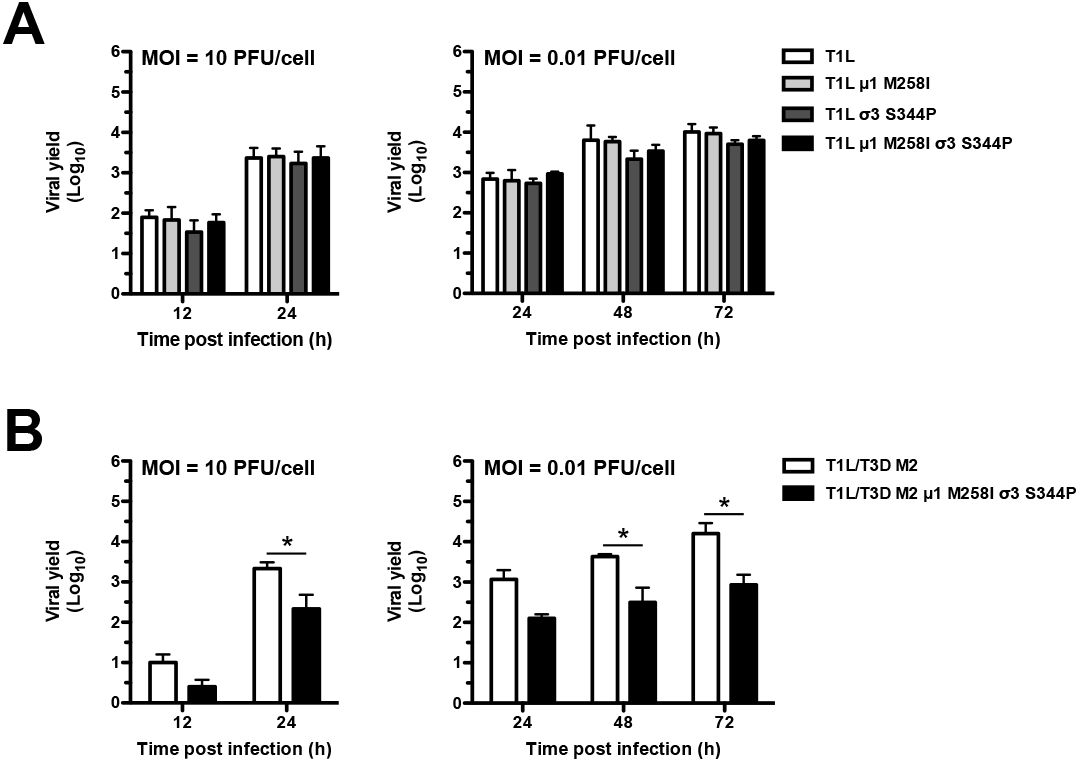
Growth profiles of T1L and T1L/T3D M2 variants. (A and B) L cell monolayers were infected with virions at an MOI of 10 PFU/cell or 0.01 PFU/cell. At the indicated times post infection, the cells were lysed and viral yield was quantified by plaque assay. The data are presented as means ± SDs. *, *P≤* 0.05 and difference in viral yield ≥ 1 log_10_ unit (n = 3 independent replicates).

### Implications for host-pathogen interactions

The σ3 protein influences cell entry (24–28), particle assembly (62–67), and environmental stability (43). Moreover, differences in the efficiency of translational shutdown map to the S4 gene segment (encodes for σ3) (68). This effect may be direct or indirect by countering protein kinase R through σ3-dsRNA interactions (69, 70). σ3 function relies on its subcellular localization and its capacity to remain unbound from μ1; the affinity between μ1-σ3 varies based on strain (71). Thus, HR mutations could impact these activities in a reassortant background Mechanistic studies are needed to dissect the relationship between the host and hyperstable reovirus.

### Conclusions

Resistance-granting mutations are tools to understand structure-function relationships, the basis (or mechanism) of inactivation, and replicative fitness. Toward this end, we selected for rare subpopulations at high temperatures (Figs. 1-2). HR strains contained two mutations within μ1-σ3. μ1 M258I was not associated with a phenotype, whereas σ3 S344P was sufficient to enhance capsid integrity and to reduce protease sensitivity (Figs. 3–4). Together, these changes impaired replicative fitness in a reassortant background (Fig. 6). This work reveals new details regarding the determinants of reovirus stability.

## ACKNOWLEDGEMENTS

We thank members of our laboratory and the Indiana University virology community for helpful suggestions. Dynamic light scattering was performed in the Indiana University Physical Biochemistry Instrumentation Facility.

Research reported in this publication was supported by the National Institute of Allergy and Infectious Diseases of the National Institutes of Health under award numbers 1R01AI110637 (to P.D.) and F32AI126643 (to A.J.S.) and by Indiana University. The content is solely the responsibility of the authors and does not necessarily represent the official views of the funders.

